# Identification of Injury-Response State Determinants in Glioblastoma Stem Cells

**DOI:** 10.1101/2025.04.07.647626

**Authors:** Fatemeh Molaei, Graham MacLeod, Shahan Haider, Aleczandria Tiffany, Frank M Oteng, Jacob M. Berman, Arianna Skirzynska, Molly S. Shoichet, Daniel Schramek, Peter B. Dirks, Stephane Angers

**Affiliations:** Donnelly Centre for Cellular and Biomolecular Research, Temerty Faculty of Medicine, Toronto, Ontario, Canada; Leslie Dan Faculty of Pharmacy, University of Toronto, Toronto, Ontario, Canada; Centre for Molecular and Systems Biology, Lunenfeld-Tanenbaum Research Institute, Mount Sinai Hospital, Toronto, Ontario, Canada; Department of Molecular Genetics, Temerty Faculty of Medicine, University of Toronto, Toronto, Ontario, Canada; Department of Biochemistry, Temerty Faculty of Medicine, University of Toronto, Toronto, Ontario, Canada; Chemical Engineering and Applied Chemistry, University of Toronto, Toronto, Ontario, Canada; Institute for Biomedical Engineering, University of Toronto, Toronto, Ontario, Canada; Department of Chemistry, University of Toronto, Toronto, Ontario Canada; Developmental and Stem Cell Biology Program and Sonia Labatt Brain Tumor Research Centre, The Hospital for Sick Children, Toronto, Ontario, Canada; Division of Neurosurgery, University of Toronto, Toronto, Ontario, Canada

## Abstract

Glioblastoma (GBM) is a common and highly lethal type of primary brain tumor in adults. Therapeutic failure is partly attributed to a fraction of Glioblastoma Stem Cells (GSCs) that show high levels of heterogeneity and plasticity. GSCs exist in a transcriptional gradient between two states: Developmental (Dev) and Injury Response (IR) in which IR-GSCs exhibit more invasive behaviors. While previous studies have identified fitness genes in GSCs, the genes required to establish and maintain the Dev and IR states remain poorly defined. To identify the regulators of the IR GSC state, we performed a phenotypic genome-wide CRISPR-Cas9 knockout (KO) screen in patient-derived GSCs based on cell surface expression of the IR marker CD44. Notably, we found that perturbations of the histone acetyltransferase *EP300* in IR *GSCs* led to decreased CD44 cell surface expression, significant downregulation of gene expression signatures associated with the IR state, and to decreased self-renewal and invasion. Furthermore, genetic targeting of *Ep300* in a mouse GBM model delayed tumor initiation and/or progression. Collectively, our results establish EP300 as a regulator of the IR state in GSCs and provide a mechanistic basis for its therapeutic targeting in GBM.

**Significance:** A genome-wide phenotypic CRISPR-Cas9 screen in a patient-derived Glioblastoma Stem Cell line identified the genes required to maintain the Injury-Response cellular state, with a focus on the histone acetyl transferase gene *EP300*. This study suggests how therapeutic targeting of cellular state could reduce the aggressiveness of GBM tumors.

## Introduction

*IDH-wild* type Glioblastoma (GBM) is the most common and aggressive form of brain tumors in adults that, despite intensive research effort, remains incurable. Genomic and transcriptomic studies in GBM tumor samples have revealed considerable intertumoral heterogeneity in bulk tumors and classified GBM tumors into three major subtypes according to their gene expression profiles: Proneural, Classical and Mesenchymal (Wang et al. 2017). However, recent single cell RNA-seq analyses revealed the detection of cells from multiple subtypes within individual tumors with the dominant subtype driving bulk tumor classification (Neftel et al. 2019). This suggests that intratumoral heterogeneity is a significant driver of the intertumoral heterogeneity observed in bulk tumors. This same study classified GBM cells into four malignant cellular states that recapitulate the transcriptomic program of neural-progenitor-like (NPC-like), oligodendrocyte-progenitor-like (OPC-like), astrocyte-like (AC-like), and mesenchymal-like (MES-like) states (Neftel et al. 2019). In addition, multiple studies have demonstrated evidence of plasticity in GBM cells, with individual clones being capable of recapitulating the heterogeneous nature of GBM tumors by giving rise to cells of multiple cellular states in xenograft models (Lan et al. 2017; Neftel et al. 2019). Therapeutic failure in GBM can largely be attributed to high levels of intratumoral heterogeneity and plasticity (Neftel et al. 2019; Louis et al. 2021).

At the root of GBM lies the Glioblastoma stem cells (GSCs) that are characterized by tumor initiation, self-renewal, and differentiation abilities (Singh et al. 2004). GSCs contribute to neovascularization, invasion, therapeutic resistance and recurrence, and are a source of intratumoral heterogeneity and plasticity (Dirkse et al. 2019). Recent single-cell and bulk RNA-seq characterization of a large panel of patient-derived cultures demonstrated that GSCs can be grouped into two major cellular states according to their gene expression profiles: Developmental (Dev) and Injury Response (IR) that underlie the previously described Proneural and Mesenchymal GBM subtypes, respectively (Richards et al. 2021). Importantly, rather than two distinct fixed subtypes, GSCs exist in a transcriptional gradient between the Dev and IR states, indicative of cellular plasticity that contributes to intratumoral heterogeneity in GBM tumors. A growing body of evidence indicates that in response to standard-of-care treatment (chemotherapy plus radiation) or exposure to cytokines, Dev/Proneural cells can transition to a more IR/Mesenchymal phenotype, which is more invasive and therapy-resistant, features that contribute to a worse prognosis (Richards et al. 2021; Hara et al. 2021; Segerman et al. 2016; Guilhamon et al. 2021; Khan et al. 2023; Bhat et al. 2013; Wang et al. 2017).

Genome-wide and targeted CRISPR-Cas9 fitness screens on panels of GSCs have revealed differences in genetic dependencies between subtypes and the existence of a gene fitness gradient between IR and Dev GSCs, functionally manifesting the transcriptional gradient (Richards et al. 2021; MacLeod et al. 2024). While these screens provide a wealth of information on functional differences between the subtypes, precisely how the transcriptional gradient between GSC subtypes is established and maintained is not fully understood, although it is likely influenced by genetic abnormalities, cell-cell interactions and clonal variation (Lan et al. 2017). Knowledge of which genes are required to maintain the IR-GSC state would greatly increase our understanding of GSC biology. In theory, if the genes or networks required for the maintenance of this more aggressive state were perturbed, malignant cells could be “locked” into a less aggressive phenotypic state and the prognosis for patients could be improved.

In this study, we performed a genome-wide flow cytometry-based CRISPR-Cas9 phenotypic screen on patient derived IR GSCs, to identify genes required to maintain the most aggressive IR-GSC state. We identified *EP300*, encoding the histone acetyltransferase p300, as a master regulator of the IR state. Genetic or pharmacological perturbations of *EP300* decreased the invasiveness and stemness properties of IR GSCs. Inhibition of *EP300* disrupted the epigenome and shifted GSCs out of the IR state, thereby inhibiting cancer progression in an *in vivo* GBM model. Taken together, our study revealed new IR phenotype regulators and established a potential therapeutic opportunity to interfere with maintenance of the IR state to reduce GBM tumorigenicity.

## Results

### Genome-wide CRISPR-Cas9 phenotypic screen identifies regulators of the Injury-response GSC state

Previous work has identified CD44 as a marker of the IR/Mes state in GBM, with higher expression in IR GSCs when compared to Dev GSCs (Richards et al. 2021; Hara et al. 2021) (Fig. 1A). Elevated expression of CD44 on the surface of IR GSCs compared to Dev GSCs was confirmed via flow cytometry in a panel of patient-derived GSC cultures (Fig. 1B-C). To identify genes required for maintenance of the IR state in GSCs, we performed a Fluorescence-Activated Cell Sorting (FACS)-based genome-wide CRISPR-Cas9 knockout (KO) phenotypic screen using CD44 cell surface expression as a fiducial marker of the IR state. To do so, we transduced a patient-derived IR GSC culture (G549) with the TKOv3 genome-wide human gRNA library (Hart et al. 2017) to generate a pool of knockout cells and isolated cells with high and low CD44 expression using FACS (Fig. 1D). A comparison of gRNA abundances using the MAGeCK algorithm in CD44-low (bottom 15%) and CD44-high (top 15%) GSC populations identified genes whose KO decreased CD44 expression levels. As expected, *CD44* was identified as the top hit enriched in the CD44-low bin (Table S1). Predictably, as CD44 is a transmembrane glycoprotein, our screen identified several genes with known functions in macromolecular/protein glycosylation (*GALNT1*, *GALNT6*) as well as *EMP3,* which has previously been shown to regulate CD44 levels at the cell surface (Fig. 1E-F) (Thornton et al. 2020; J. Zhang et al. 2022). In addition, we identified *JUN* and *FOSL1,* which have previously been reported as regulators of the mesenchymal signatures in GBM (Fig. 1E) (Marques et al. 2021; Lv et al. 2022). Notably, several additional transcription factors (TFs), such as *SRF* and *KLF6,* and chromatin remodeling genes including *EP300* and *SETD1B* were also enriched in the CD44-low population, suggesting their requirement in sustaining transcriptional programs defining the IR state.

**Figure 1.**
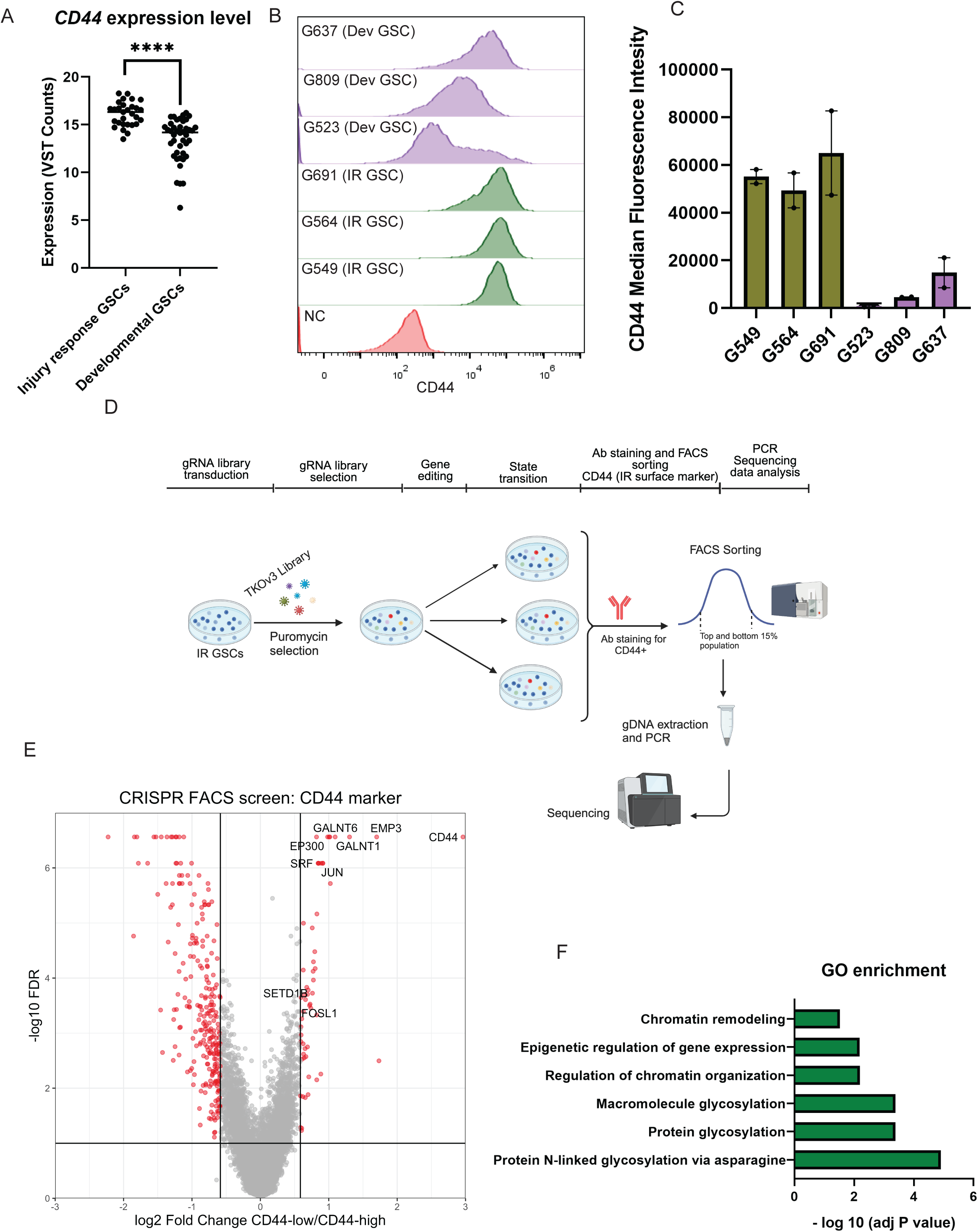
CRISPR-Cas9 phenotypic screen identifies genes required to maintain the IR GSCs state. A) *CD44* mRNA expression in injury response (IR) (n=29) vs developmental (Dev)(n=42) patient derived GSC cultures. B) Flow cytometry analysis of CD44 cell surface expression in a panel of patient-derived IR and Dev GSC cultures (N=2). Mean Fluorescence Intensity (MFI). HEK293 cells were used as negative control (NC) C) Quantification of CD44 expressed as median fluorescent intensity (MFI) in IR and Dev GSCs (N=2) from panel (B). D) Schematic of genome-wide phenotypic CRISPR screens for identification of genes required to maintain the IR GSCs state. E) Volcano plot of genes enriched in CD44 “low” population compared to CD44 “high” population. p-values from genome-wide CRISPR screen calculated by the MAGeCK algorithm. Selected hits, including previously identified regulators of IR/Mes state (*JUN, FOSL1*), CD44 glycosylation/trafficking (*GALNT6, GALNT6, EMP3*) and chromatin remodelling (*EP300,SRF, SETD1*) are highlighted. F) Selected Gene Ontology (GO) for hits enriched in CD44 “low” population compared to CD44 “high” population (FDR less than 0.2)

### Loss *of EP300* compromises self-renewal and tumor initiation of IR GSCs

Transition between cell states requires epigenetic reprogramming driven by alterations in chromatin accessibility, a process in which histone acetylation plays an important role (Reik, Dean, and Walter 2001). EP300 is a bromodomain histone acetyltransferase, which, along with cyclic AMP (cAMP) response element binding protein binding protein (CREBBP), is responsible for histone H3 lysine 27 acetylation (H3K27Ac) at super-enhancer and promoter regions and is a mark associated with active gene transcription (Dancy and Cole 2015; Ogryzko et al. 1996). EP300 function affects the expression of genes crucial for cell cycle regulation, apoptosis, and DNA repair, thereby shaping the cancer cell phenotype (Q. Chen et al. 2022). Additionally, EP300 interacts with various transcription factors and acts as a transcriptional coactivator (Ogryzko et al. 1996).

Knockout of *EP300* using two different gRNAs in two IR GSCs (G549 and G411) resulted in decreased CD44 surface expression when compared to control (Fig. 2A and C, S1A and C). Treatment with A-485, a selective inhibitor of EP300 histone acetyltransferase activity, showed decreased levels of CD44 in two IR GSCs (G549 and G411) when compared to cells treated with DMSO (Fig. 2B-C, S1B-C) (Lasko et al. 2017). Collectively, these findings verify our screen results and confirm that EP300 regulates surface expression of CD44 in IR GSCs.

**Figure 2.**
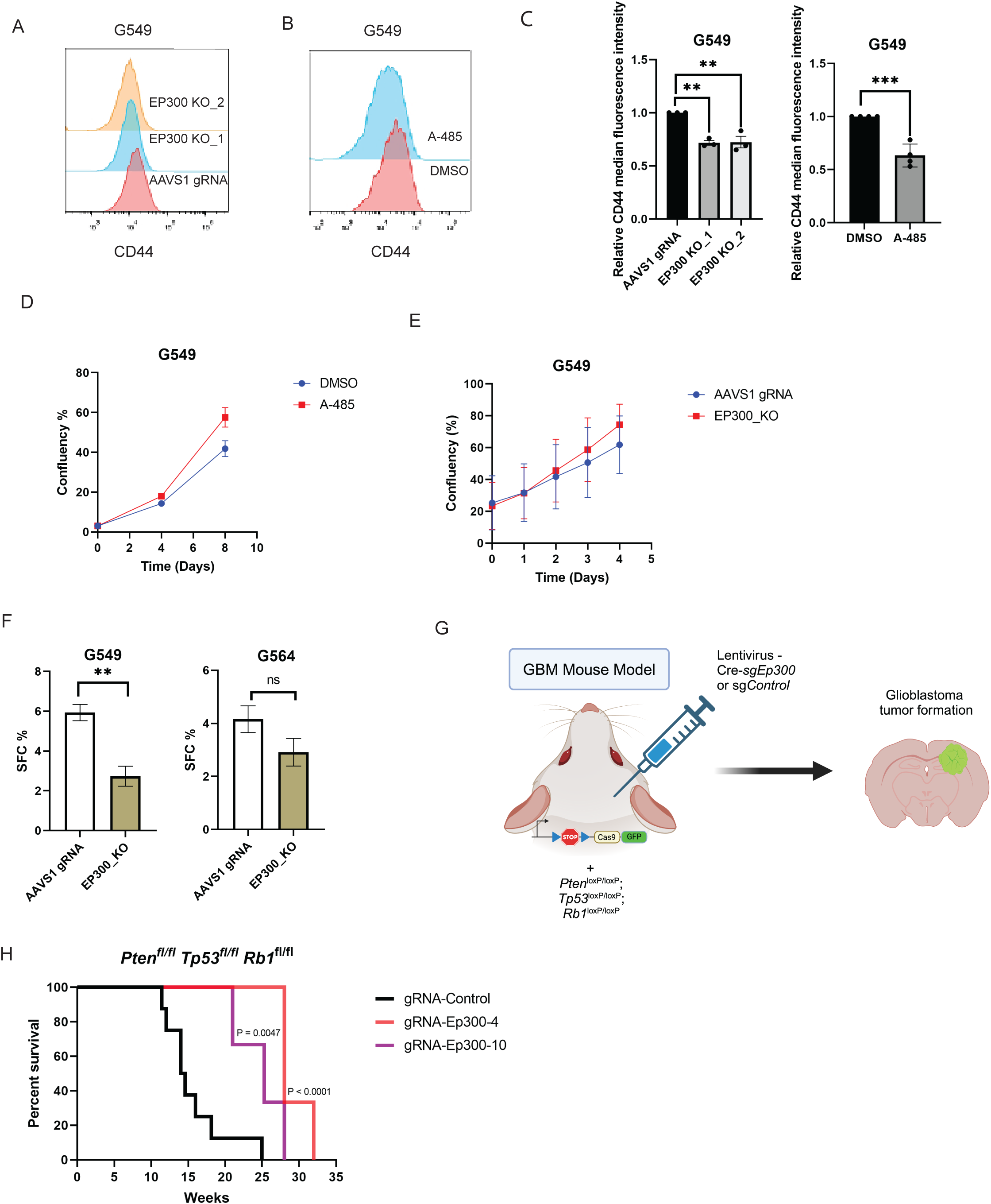
*EP300* is required for self-renewal of IR GSCs and tumor initiation and/or progression *in vivo*. A) Representative Flow cytometry experiments (3 independent experiments) measuring CD44 cell surface expression in IR GSC (G549) transduced with gRNA targeting *AAVS1* (control) or *EP300* (2 gRNAs). B) Representative Flow cytometry experiment (3 independent experiments) measuring CD44 surface expression following treatment of IR GSCs (G549) for 7 days with DMSO or A-485. C) Bar graph representing the quantification of CD44 cell surface expression (expressed as median fluorescent intensity relative to the control) in panel (A) and (B) (n=3). Error bars: mean ± SEM,* p<0.05 ** p<0.01. D) Quantification of IR GSC (G549) proliferation following treatment with control (DMSO) or with 400nM EP300 inhibitor (A-485) (n=3) using the Incucyte Live Cell Imaging system. Error bars: mean ± SEM. E) Quantification of IR GSC (G549) proliferation following control (*AAVS1 gRNA)* or *EP300* knockout (*EP300_KO)* (n=3). Error bars: mean ± SEM. F) *In vitro* limiting dilution assay (LDA) of GSCs culture. Sphere forming Capacity (SFC) of the indicated GSC cultures expressing gRNA targeting *AAVS1* (control) or *EP300*. Error bars: estimated frequency +95% C, ** p<0.01. G) Schematic illustrating the GBM mouse model (Created with BioRender.com). Mice carrying LSL-Cas9-GFP and triple floxed *Tp53*, *Pten* and *Rb1* alleles were intracranially injected with lentivirus carrying Cre-recombinase and gRNAs targeting either control or *Ep300*. I) KaplanMeier curve for *Tp53/Pten/Rb1* triple knockout GBM mouse model in tumors expressing the indicated gRNAs. Indicated p-value was calculated using the log-rank (Mantel-Cox) test.

Next, we assessed the effect of EP300 perturbations on proliferation of IR GSCs. We observed that the proliferation rate of IR GSCs was not significantly affected by *EP300* KO or chemical inhibition when compared to controls (Fig. 2D-E, S1D-E). This is consistent with previous CRISPR fitness screens performed by our lab where *EP300* was not identified as a genetic vulnerability in IR GSCs (MacLeod et al. 2019, 2024) (Fig. S1F). One of the features of the CD44 phenotypic screen was the ability to identify regulators of cell state whose KO may not affect cell growth or survival but that affect other phenotypes related to the IR state. Importantly, we observed that the sphere-forming capacity (a surrogate for self-renewal capacity) of IR GSCs was decreased upon *EP300* KO (Fig. 2F, S1G). This finding suggests EP300 is a regulator of stemness in IR GSCs as opposed to proliferation. Increased resistance to chemotherapy is a feature often associated with mesenchymal/IR GSCs, therefore, we evaluated the effect of *EP300* KO on temozolomide (TMZ) sensitivity. We observed that *EP300* KO did not change the sensitivity of IR GSCs to TMZ (Fig. S1H). This aligns with our previous findings, where *EP300* was not among the genes identified to modulate GSCs sensitivity to TMZ treatment (MacLeod et al. 2019).

To investigate the role of *EP300* in GBM tumor initiation and progression, we tested the requirement of *Ep300* in a mouse model of GBM consisting of disruption of 3 common GBM tumor suppressor genes *Tp53, Pten* and *Rb1* (Chow et al. 2011; MacLeod et al. 2024). Mice carrying floxed alleles of *Tp53* (*Tp53*^fl/fl^), *Pten* (*Pten*^fl/fl^) and *Rb1* (*Rb1*^fl/fl^) as well as LSL-Cas9-GFP were subjected to stereotactic injection at P0 to deliver lentiviral particles expressing Cre and either *Ep300*-targeting or control gRNAs and transduce the mouse neural stem cells residing in the lateral subventricular zone (Fig. 2G). Kaplan Mayer survival analysis showed that genetic ablation of *Ep300,* using two independent gRNAs, significantly prolonged the survival of mice when compared to control gRNA-injected animals (Fig. 2H). While the virus induced efficient *Ep300* gene editing in MEF and NSC cell lines, all tumors that grew out in mice showed no evidence of gene editing, indicating that they are likely “escapers” (Fig. S2). Together, this data shows that *Ep300* is required for tumor initiation and/or progression.

### EP300 is required to maintain the invasive behavior of IR GSCs

Based on Single Cell ATAC-seq profiling, GSCs have been characterized into three states: Constructive, Reactive, or Invasive (Guilhamon et al. 2021). Invasive and Reactive GSC cultures have IR expression signatures, thus likely representing further functional stratification of the IR subtype, while the Constructive GSCs align with Dev signatures. One of the main features of Invasive/IR GSCs is the ability to invade surrounding tissues and contribute to the aggressiveness of tumors (Guilhamon et al., 2021). To investigate the requirement of *EP300* for the invasive behavior of IR GSCs, we used a hyaluronic acid-based hydrogel to mimic tumor ECM and microenvironment to perform invasion assays (Smith et al. 2023) (Fig. 3A). *EP300* knockout decreased both the depth of invasion and percentage of invasive cells in two independent IR GSC cultures (G411 and G549) (Fig. 3B-E). Supporting these results, inhibition of EP300 using A-485 also significantly decreased both the the percentage and depth of invasion (Fig. 3B-E). We conclude that EP300 regulates the invasive phenotype of IR GSCs.

**Figure 3.**
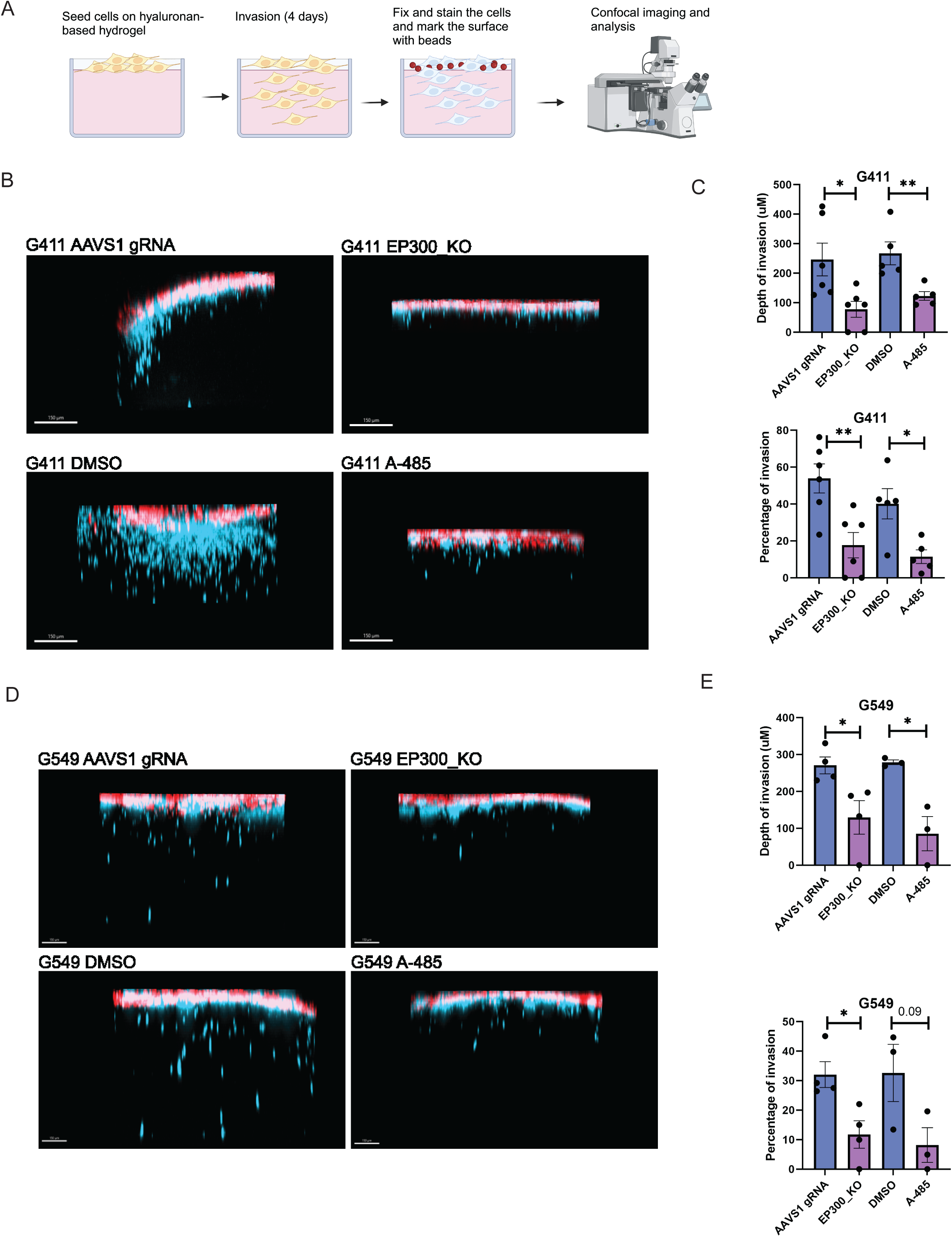
EP300 is required for invasiveness of IR GSCs. A) Schematic of the invasion assay. GSCs were seeded on top of the hydrogel, and allowed to invade for 4 days. Subsequently, cells were fixed and the surface of the gel was marked using beads for further analysis. B) Representative compressed z-stack confocal images of G411 IR GSCsc seeded and invading the hydrogel (Cells are labelled in blue and beads are in red). Cells are either control, knockout for *EP300* or pre-treated with the EP300 inhibitor A-485 for 7 days at 400 nM. C) Depth of invasion and percentage of invading cells upon *EP300* KO or chemical inhibition of EP300 (pre-treatment with A-485 for 7 days at 400 nM) in G411 IR GSCs decreased significantly when compared to control. (*n* = 3, Error bars: mean ± SEM, Unpaired T-test, \**p* < 0.05, \*\**p* < 0.01) D) Representative compressed z-stack confocal images of G549 IR GSCs seeded and invading the hydrogel (Cells are labelled in blue and beads are in red). Cells are either control, knockout for *EP300* or pre-treated with the EP300 inhibitor A-485 for 7 days at 400 nM. E) Depth of invasion and percentage of invading cells upon *EP300* KO or chemical inhibition of EP300 (pre-treatment with A-485 for 7 days at 400 nM) in G549 IR GSCs decreased significantly when compared to control. (*n* = 3, Error bars: mean ± SEM, Unpaired T-test, \**p* < 0.05, \*\**p* < 0.01).

### *EP300* activity is required to maintain the Injury Response transcriptional signature

GSCs in the IR state are characterized by high expression of mesenchymal cell-related genes and pathways associated with immune response, inflammation, wound healing, extracellular matrix and cell adhesion (Richards et al. 2021). Since we observed that *EP300* perturbations reduce several key phenotypes associated with IR GSCs (CD44 expression, self-renewal, tumor initiation and invasion), we next investigated the requirement of *EP300* to maintain the transcriptional signature of IR GSCs. RNA-seq and differential expression analysis revealed significant downregulation of multiple genes associated with the IR state, including *ITGAX, CD44, SERPINE1,* and *SERPINA1* in *EP300* KO samples when compared to control cells (Fig. 4A, S3A). Gene Ontology (GO) enrichment analysis of downregulated genes revealed overrepresentation of major pathways associated with the IR state, including ECM, wound healing and inflammation response (Fig. 4B-D), while differentially upregulated genes were enriched for cellular and tissue development pathways (Fig. S3B). Gene set enrichment analysis (GSEA) on RNA-seq results evidenced the downregulation of Mesenchymal GBM and Hallmark Epithelial-to-Mesenchymal (EMT) signatures upon *EP300* KO (FDR < 0.05) (Fig. 4E, S3C) (Phillips et al. 2006; Wang et al. 2017). Furthermore, based on the GSEA, we observed significant loss of inflammation pathways (SCHMITT TNF, and HMG) that are required to induce PN to Mes transition and to induce the Mes signature in GBM cells (Schmitt et al. 2021) (Fig. S3C). Together, our results suggest that *EP300* regulates the IR phenotype and is required to maintain the IR transcriptional program.

**Figure 4.**
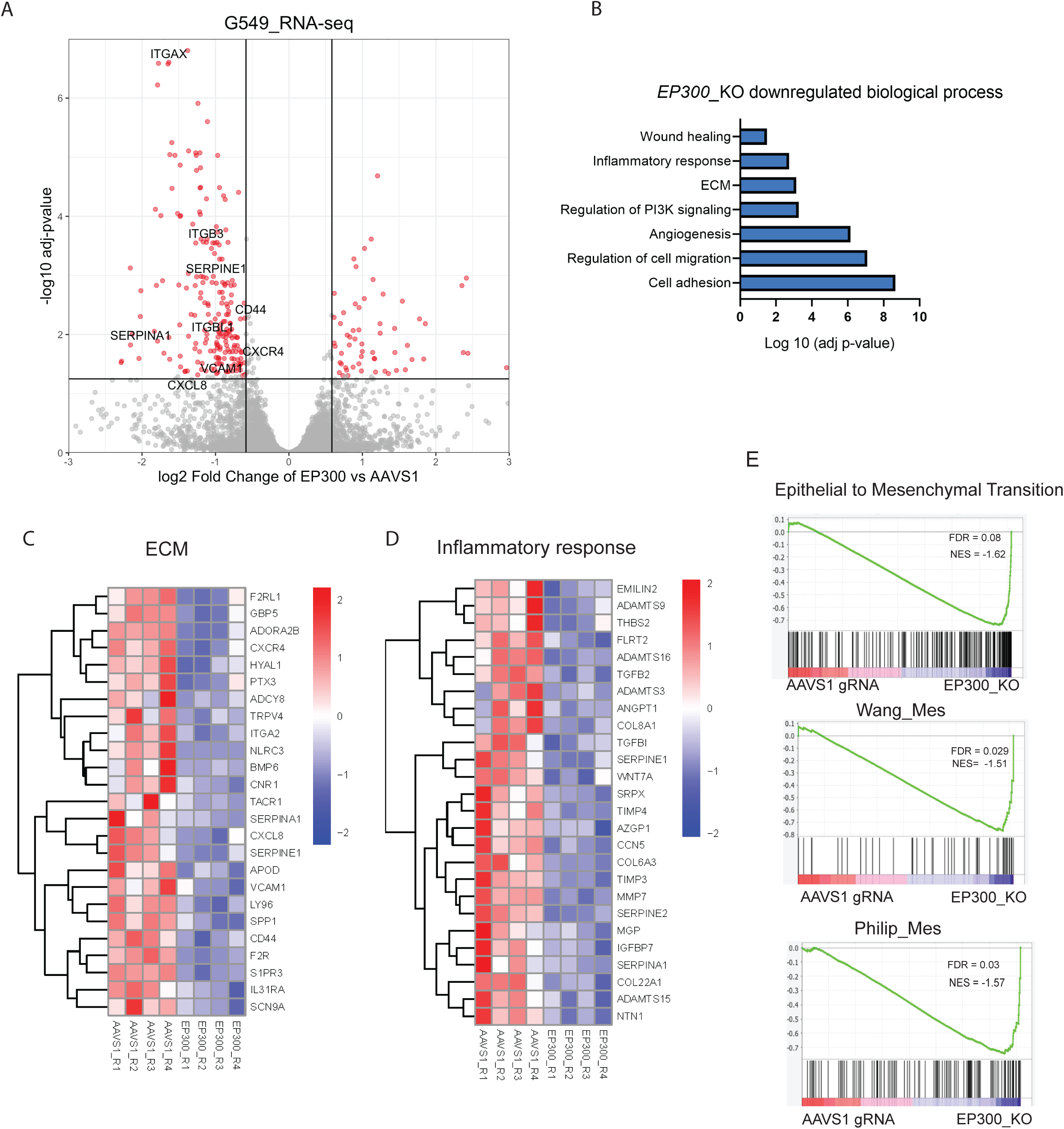
EP300 is required to maintain IR response gene signature in GSCs. A) Volcano plot depicting differentially expressed genes in G549 IR GSCs knockout for *EP300* vs control (*AAVS1* gRNA). N=4. B) Selected Gene Ontology (GO) biological process differentially downregulated in IR GSCs knockout for *EP300* (n=4). C) Heatmap comparing average expression of genes associated with the Extracellular Matrix GO term (GO:0031012) in control or *EP300* knockout IR GSCs. D) Heatmap of average expression of genes in Inflammatory Response GO term (GO:0006954) in *EP300* KO samples vs control (*AAVS1* targeting gRNA). E) Representative plots of gene set enrichment in differentially expressed genes comparing *EP300* KO samples vs control (*AAVS1* targeting gRNA).

To further investigate the role of EP300 in IR GSCs, we performed chromatin profiling CUT&RUN experiments using a H3K27Ac antibody to compare *EP300* KO and control (*AAVS1* gRNA) IR GSCs. As expected, given the HAT function of EP300, we observed a global reduction of H3K27Ac signal upon *EP300* KO (Fig. S4A). In *EP300* KO cells, a clear reduction of H3K27Ac signal around the TSS was observed for genes whose expression was downregulated upon *EP300* KO (Fig. 5A). Our results therefore confirm that EP300 regulates transcriptional programs in IR GSCs via the H3K27Ac epigenetic modification. Next, to test whether EP300 modulates the IR state through a direct or indirect mechanism, we performed CUT&RUN in IR GSCs using an EP300 antibody and identified 2800 regions occupied by EP300 (SEACR peakCalling using 0.001 q value cutoff) (Fig. 5B). Importantly, we observed a strong H3K27Ac signal reduction around EP300 binding sites in *EP300* KO GSCs suggestive of direct epigenetic regulation at these loci (Fig. 5C). Importantly, this includes several IR signature genes found to be regulated by EP300 in our RNA-seq experiment (e.g.: *CD44*, *CXCR4* and *SERPINE1*) (Fig. 5D). We conclude that EP300 is directly involved in the regulation of the IR transcriptional state.

**Figure 5.**
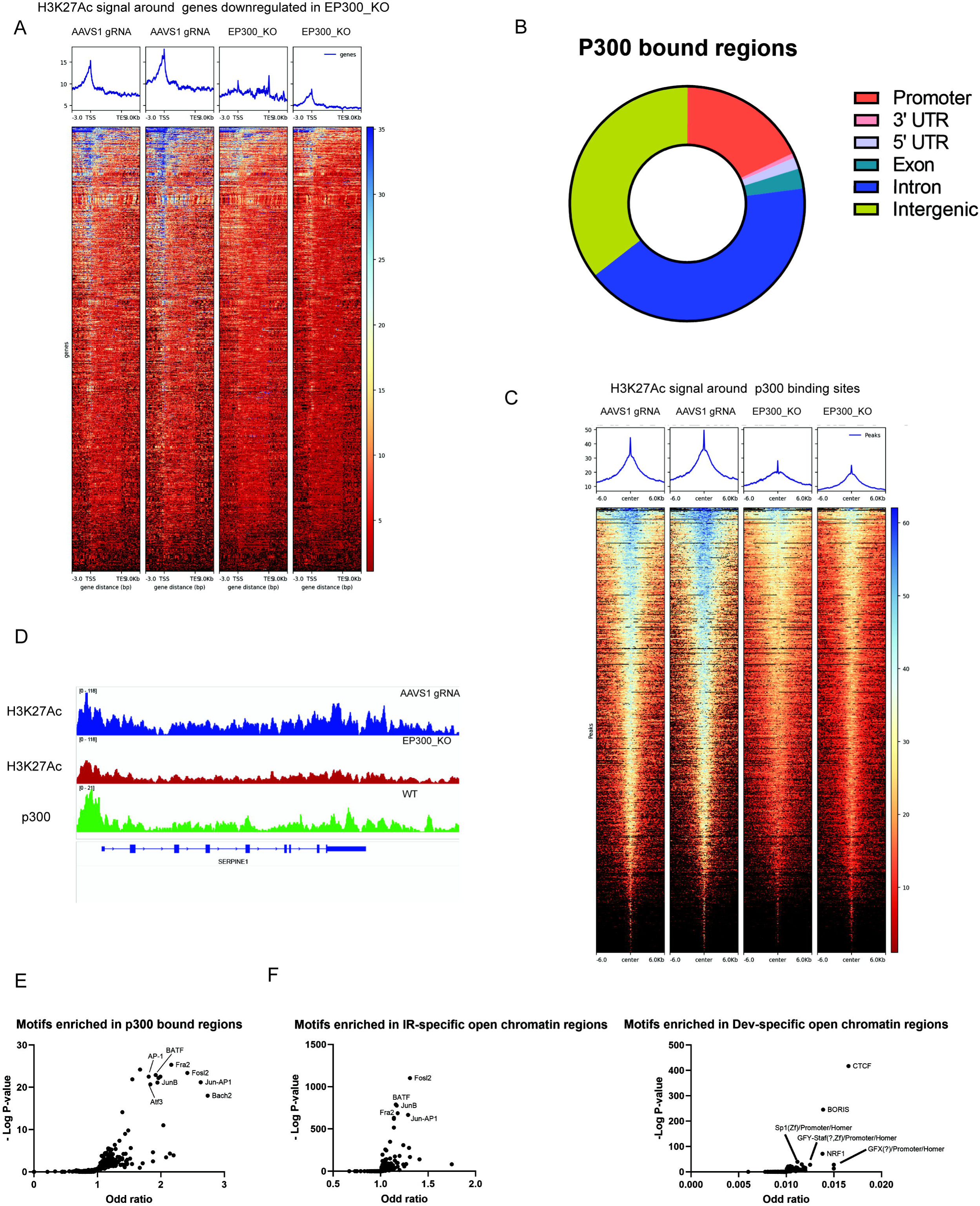
Regulation of H3K27Ac by EP300 modulates the IR signature. A) Heatmap of H3K27Ac CUT&RUN signal in control (AAVS1 targeting gRNA) and *EP300* KO IR GSCs (G549) around regions of genes found to be transcriptionally down-regulated upon *EP300* KO. B) Doughnut plot displaying genomic distribution of EP300-binding sites. C) Heatmap of H3K27Ac CUT&RUN signal in control (AAVS1 targeting gRNA) and *EP300* KO IR GSCs (G549) around EP300 bound regions D) Representative genome browser image at the *SERPINE1* locus displaying H3K27Ac signal in control (*AAVS1* targeting gRNA) (Blue) and *EP300* KO (Red) IR GSCs (G549), or EP300 binding signal (green) in WT IR GSCs (G549) from CUT&RUN experiments. E) Dot plot displaying TF motifs significantly enriched in EP300 binding sites F) Dot plot displaying TF motifs significantly enriched in IR-GSC and Dev-GSC open chromatin regions identified by ATAC-seq (Guilhamon et al).

Analysis of EP300 bound regions revealed enrichment of JUN-AP1, FRA-1 and FRA-2 family binding motifs (Fig. 5E). These transcription factor families have previously been shown to drive expression of the mesenchymal GBM signature and are consistent with our screen results revealing members of the JUN and FOS families as IR state regulators (Marques et al. 2021; Lv et al. 2022). We further assessed the motifs enriched within enhancer regions of the H3K27Ac CUT&RUN dataset and identified significant enrichment of JUN and FOS family binding motifs in control samples, a signal that was lost in *EP300* KO cells (Fig. S4B). To investigate the importance of these EP300-regulated regions in modulating the IR phenotype across a larger set of GSCs, we analyzed previous genome-wide ATAC-seq data and the open chromatin regions that were enriched in IR (19 independent patient cultures) and Dev (7 independent patient cultures) GSCs (Guilhamon et al. 2021). We observed that the motifs enriched in EP300-bound regions (JUN and FOS family) overlapped with motifs enriched in the IR GSC open chromatin regions but not Dev GSC open chromatin regions (Fig. 5F). We conclude that EP300 modulates the IR GSC phenotype through regulation of H3K27Ac levels and IR-specific genes transcriptional programs in cooperation with TFs that are required to maintain the state.

## Discussion

Given the heterogeneity and plasticity of GBM tumors, we now understand why targeted single-agent therapies have failed to have a significant clinical impact. Further studies that elucidate how cellular heterogeneity is established and maintained in GSCs, and better characterization of the plasticity between cellular states are needed to identify new therapeutic targets or alternative strategies that could more effectively harness the malignant cells that survive front-line therapy.

Within the Developmental-Injury Response axis in GSCs, considerable heterogeneity exists at genetic, epigenetic, transcriptional, and functional levels (Richards et al. 2021; Guilhamon et al. 2021; MacLeod et al. 2024). Dev GSCs are able to convert into IR state through different mechanisms, including exposure to radiation or secreted chemokines and activation of certain TFs (E.g: *STAT3*, *TAZ, FOSL1, JUN*), giving rise to a more heterogeneous population of cells (Richards et al. 2021; Bhat et al. 2011, 2013; Lau et al. 2015; Lv et al. 2022; Marques et al. 2021). However, the mechanisms underlying maintenance of the IR state and IR-to-Dev transition are not fully understood. The CRISPR-Cas9 phenotypic screen presented in this study was designed to identify regulators of the IR state based on a well-established cell surface marker, CD44. Among the identified candidate IR state regulators were *FOSL1* and *JUN,* which were previously shown to maintain the mesenchymal signature in GBM tumors. Interestingly, the screen identified the HAT and transcriptional coactivator *EP300* as required for CD44 surface expression raising the hypothesis that it is essential to maintain the IR state in GSCs. Epigenetic regulation and chromatin remodeler genes play a critical role in cell state transitions as well as maintenance of cell states (Suvà, Riggi, and Benstein 2013). In GBM, EP300 is required for radiation-induced state transition that contributes to a more aggressive and therapy-resistant phenotype (Muthukrishnan et al. 2022). In the present study, we showed that *EP300 KO* leads to loss of the IR gene signature as well as loss of H3K27Ac signals around the genes downregulated upon *EP300* KO, which reinforces the importance of EP300 in directly governing the IR state. Binding motifs for TFs (JUN/FOS) known to drive IR/Mes signatures were significantly enriched at EP300 binding sites. Furthermore, these same motifs were enriched at active enhancers in IR GSCs, a signal that was lost upon *EP300* KO. Our finding shows that EP300 is needed to maintain the transcriptional network of IR-related genes through two possible processes: 1) Interaction with TF facilitating their binding to the DNA to activate the IR program and 2) through maintaining H3K27Ac within enhancer regions of IR-related genes.

Mesenchymal cells typically express genes related to epithelial-mesenchymal transition (EMT), a process associated with tumor progression and invasion (Chanoch-Myers et al. 2022). EP300 was previously linked to EMT in several cancers and contributes to tumor progression. In Triple negative breast cancer, EP300 regulates the cancer stem cell phenotype and contributes to tumor invasive phenotype (C.-H. Li et al. 2023). In hepatocellular carcinoma, EP300 induces EMT and invasion of tumor cells through activation of Elk1-aPKC-ι signaling (Ma et al. 2020). Furthermore, EP300 induces invasion, migration and EMT in oral squamous cell carcinoma through activation of a TGF-βRII/EP300/Smad4 cascade (D. Zhang et al. 2023). In another study that demonstrated Smad3/TGF-**β**1 mediated regulation of a mesenchymal GSC signature, EP300 was found to interact with Smad3 in mesenchymal but not proneural GSCs (Fan et al. 2021). Moreover, it was shown that inhibition of EP300 in these cells reduces self-renewal (Fan et al. 2021). In this study, we directly demonstrate the importance of EP300 function in maintaining IR gene signatures and validate that this signature includes genes important for EMT.

ATAC-seq on GBM tumor cells revealed the presence of three GSC states based on chromatin accessibility profiles (Constructive, Reactive and Invasive). GSC cultures exhibiting an IR transcriptional signature are split between the invasive and reactive states, suggestive of further functional heterogeneity within the IR subtype. The Invasive GSCs have an increased capacity to invade the surrounding tissues and contribute to the poor survival and aggressive phenotypes (Smith et al. 2023; Guilhamon et al. 2021). In our study, we showed that EP300 has an important role in invasion and self-renewal ability of IR-GSCs. This finding indicates that targeting EP300 to drive cells out of the IR state, could potentially alleviate some of the most problematic features associated with these cells including their invasive behavior and increased self-renewal properties.

Importantly, our study showed that KO of *Ep300* in a mouse GBM model delayed the development of tumors suggesting that *EP300* plays a critical role in tumor initiation and/or progression. These results provide a mechanistic basis for testing the efficacy of existing EP300 inhibitors in human studies with the notion that EP300 activity may be especially important in the context of GBM tumor dominantly expressing a mesenchymal signature.

In this study we focused on identifying genes important for the IR state with the hypothesis that strategies directed to drive GSCs out of this aggressive state would interfere with tumor progression. Our findings showing that the epigenetic regulator EP300 governs the expression of several genes differentially expressed between the IR and Dev subtypes highlight EP300 as a master regulator of the IR state. The results showing that inhibition of EP300 function interfered with GSC IR phenotypes (invasion, self-renewal) without perturbing cell proliferation, suggest that targeting cell state may be a viable therapeutic opportunity beyond focusing on genes important for growth. Importantly EP300 inhibitors such as Pocenbrodib (NCT06785636), Inobrodib (Nicosia et al. 2023) (NCT04068597, NCT03568656), EP31670 (NCT05488548) and OPN-6602 (NCT06433947) (Matusow et al. 2024))(De Bono et al. 2019) are all in various stages of clinical trials for cancer treatment and should be evaluated for efficacy in GBM.

## Supplementary Figures

**Supplementary Figure 1.**
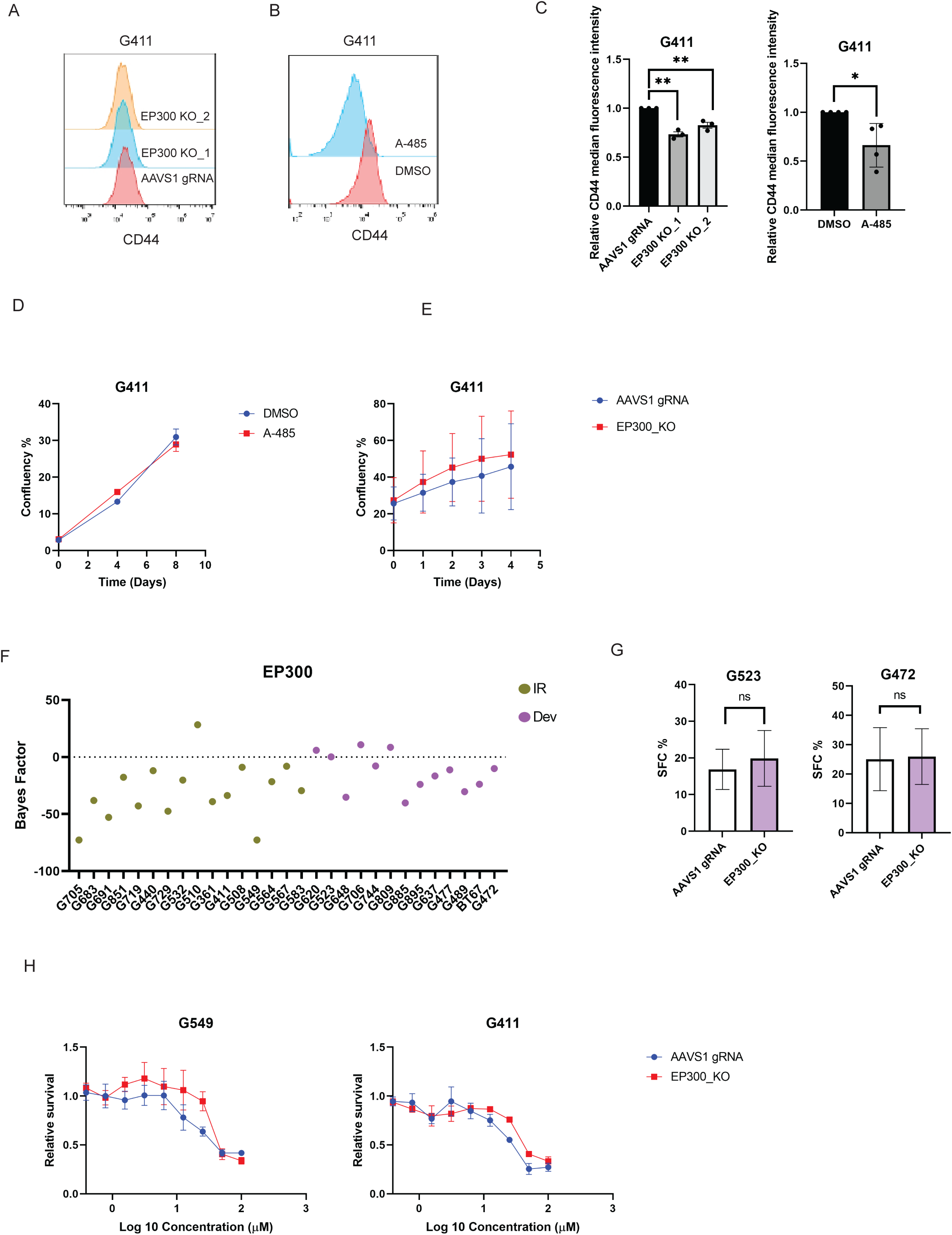
A) Representative Flow cytometry analysis of CD44 surface expression in *AAVS1* control and *EP300* KOs (2 gRNAs) in IR GSC (G411). B) Representative Flow cytometry analysis of CD44 surface expression in DMSO and A-485 treatment conditions in IR GSCs (G411). C) Bar graph representing the quantification of CD44 cell surface expression for panels (A) and (B). Expressed as a fraction of control. (n=3). Error bars: mean ± SEM,* p<0.05 ** p<0.01. D) Quantification of IR GSC (G411) proliferation following treatment with DMSO or A-485 (n=3). Error bars, mean ± SEM. E) Quantification of IR GSC (G411) proliferation following control (*AAVS1 gRNA)* or *EP300_KO* (n=3). Error bars, mean ± SEM. F) Bayes Factor (BF) scores for *EP300* from GBM5K screens in a panel of patient derived GSCs. Horizontal line indicates zero BF and any BF factor below zero is considered as non-essential. G) *In vitro* limiting dilution assay (LDA) of Dev GSCs culture. Sphere forming Capacity (SFC) for control (*AAVS1 gRNA)* and *EP300* KO in 3D cultures. Error bars, estimated frequency +95% C, ** p<0.01. H) Dose response curves of two IR GSCs comparing control (*AAVS1* gRNA) and *EP300* KO cells treated with TMZ. Error bars indicate sem from n=3 biological replicates per line.

**Supplementary Figure 2.**
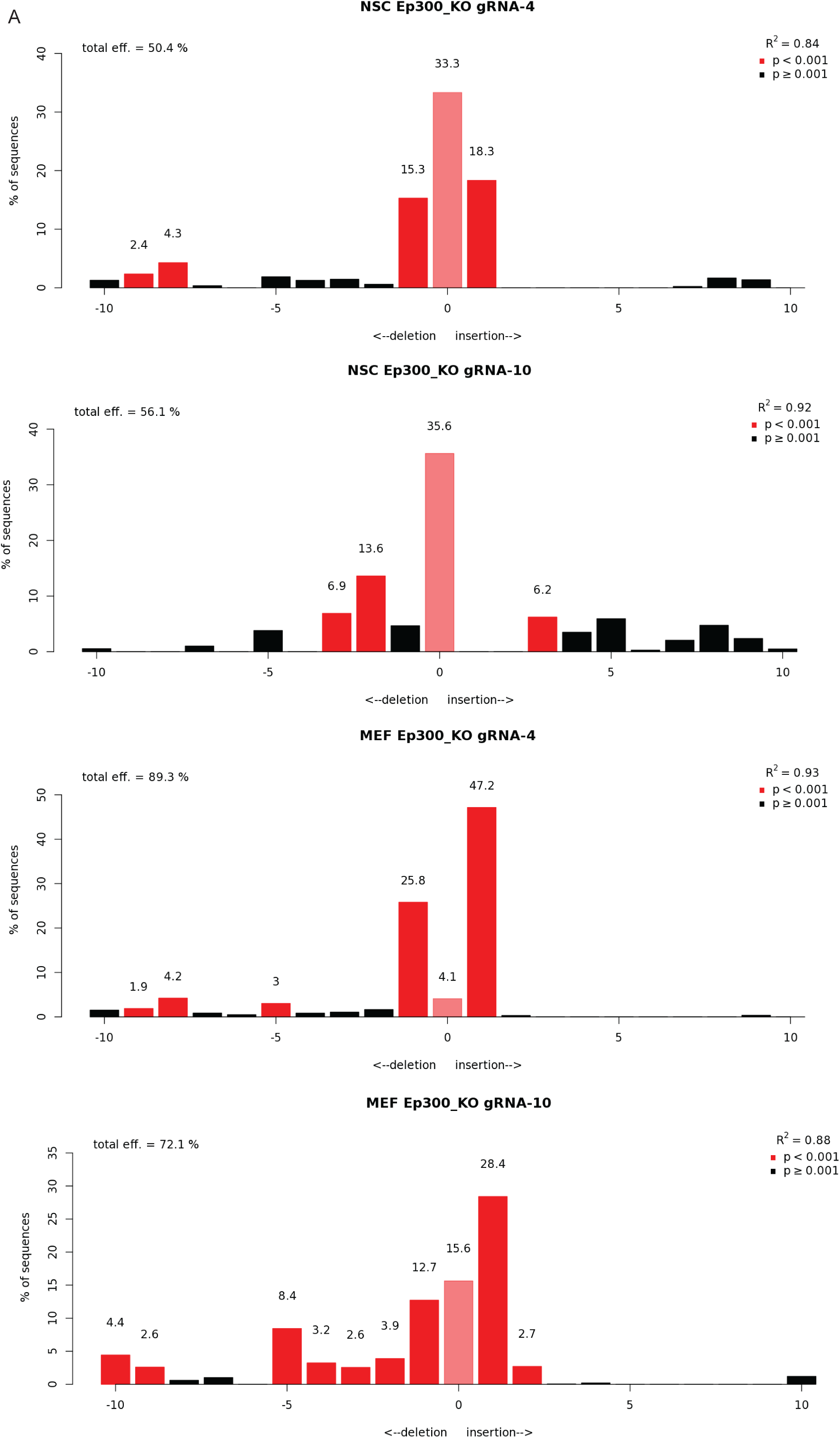
A) TIDE analysis of *Ep300* gRNA #4 and #10 editing efficiency in NSC and MEF cells.

**Supplementary Figure 3.**
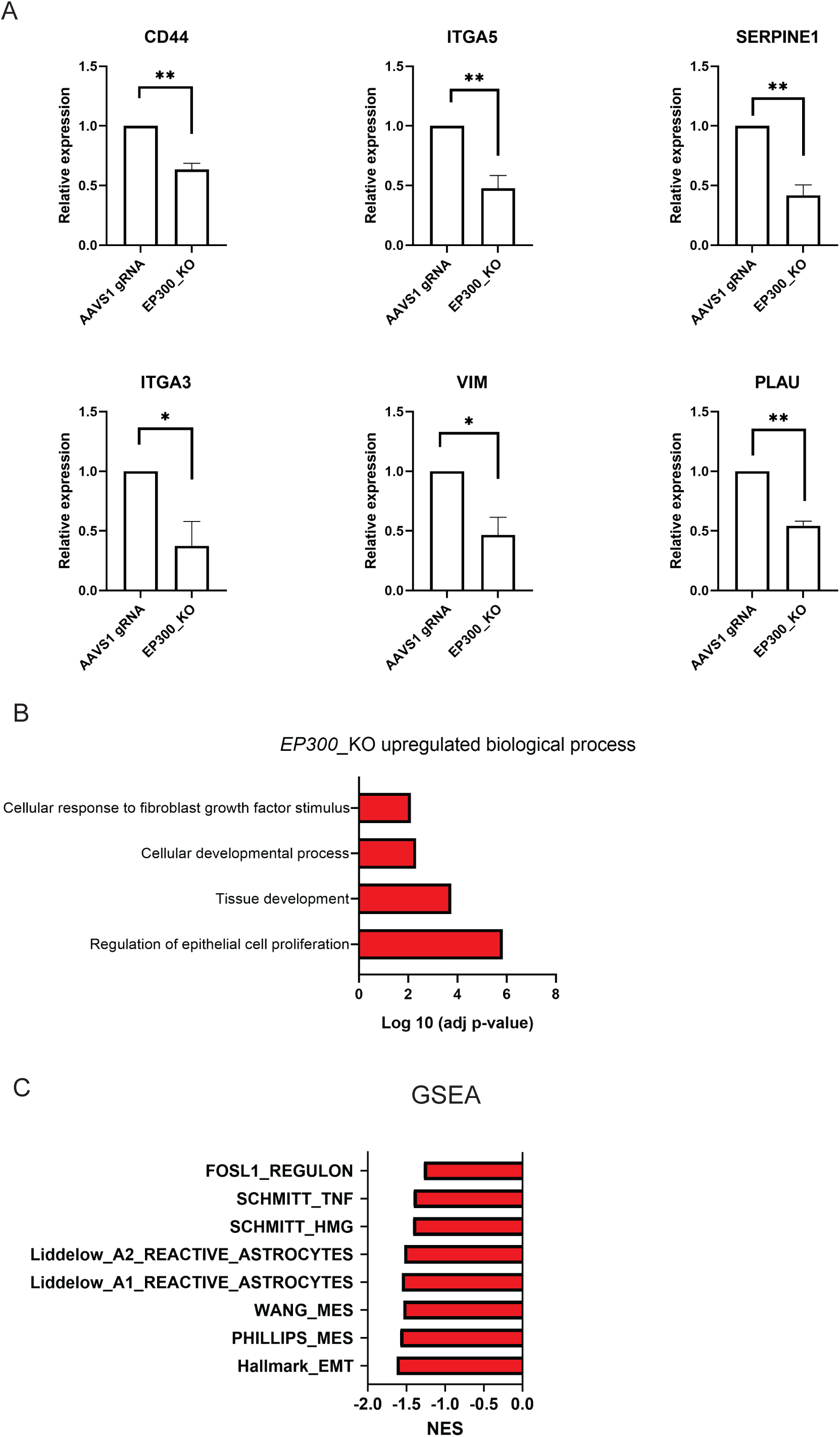
A) qPCR analysis of *CD44, SERPINE1, VIM, PLAU, ITGA3, and ITGA5* mRNA levels in G549 GSCs upon transduction with control (*AAVS1* gRNA) or *EP300* targeting gRNA (N=3). Two tailed unpaired t-test ( \**p* < 0.05, \*\**p* < 0.01). Error bars Error bars, mean ± SEM. B) Selected enriched Gene Ontology (GO) biological process in genes upregulated in the *EP300*_KO samples when compared to control *AAVS1* gRNA) (n=4). C) Gene set Enrichment Analysis (GSEA) of G549 GSCs RNA-seq comparing control (*AAVS1* gRNA) and *EP300* KO (FDR < 0.1).

**Supplementary Figure 4.**
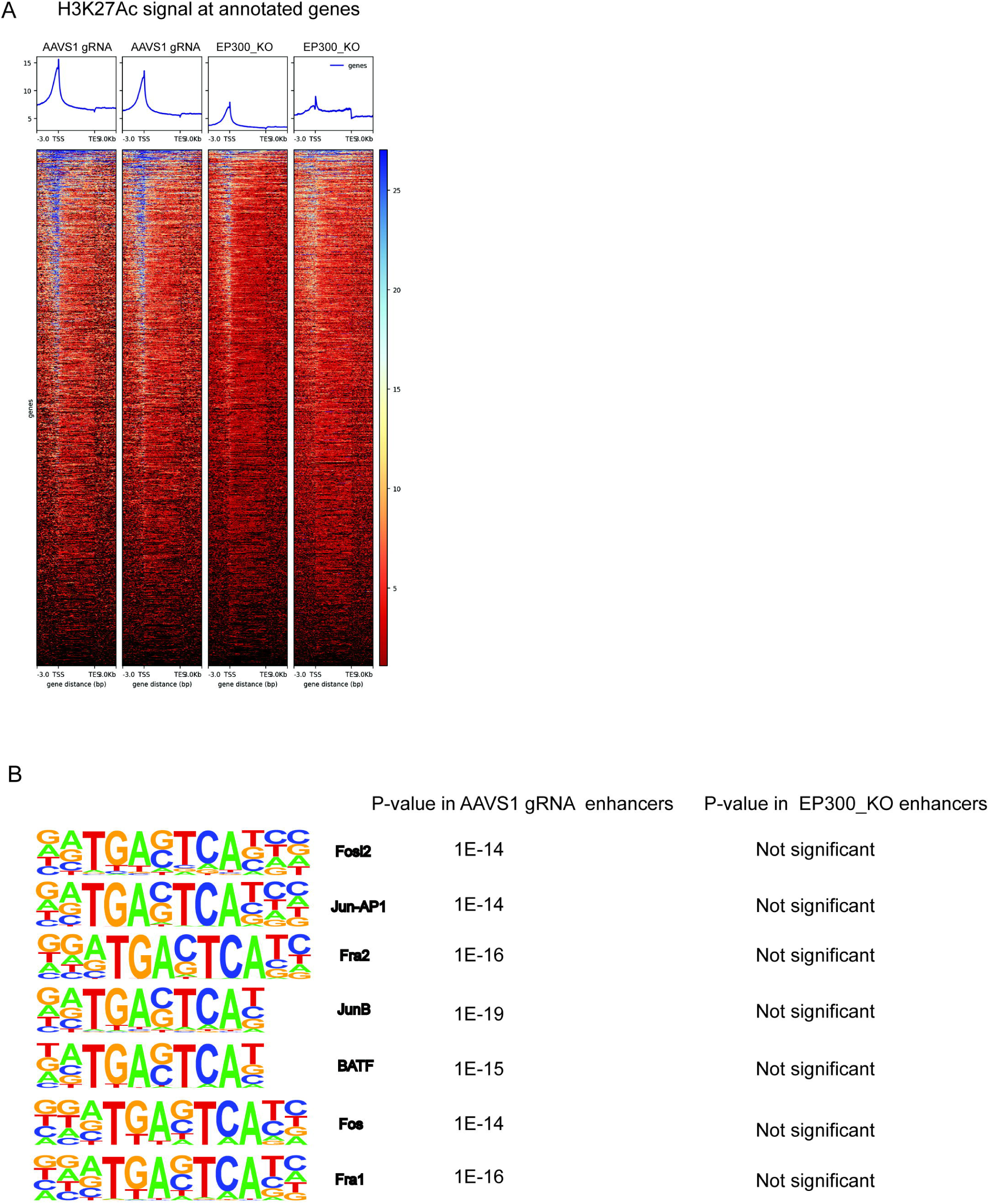
A) Heatmap of H3K27Ac CUT&RUN signals in control (*AAVS1* gRNA) and *EP300* KO G549 cells at all annotated genes (n=2 replicates per group). B) Representative motifs enrichment at enhancers in control (AAVS1 *gRNA)* G549 cells, compared to all samples (Compared to consensus peak file that includes both control and *EP300*_KO enhancer peaks).

## Methods

### Cell culture

Glioblastoma stem cell cultures were previously derived from brain tissue collected with informed consent from patients (Richards et al. 2021; Guilhamon et al. 2021; Rajakulendran et al. 2019). All experimental procedures of this study were conducted in accordance with the Research Ethics Boards at The Hospital for Sick Children (Toronto, Canada) and the University of Toronto. Glioblastoma stem cell lines were grown adherently on Poly-L ornithine (Sigma Aldrich) and laminin (Sigma Aldrich) coated plates. Cells were cultured in Neurocult NS-A Basal media (StemCell Technologies) supplemented with 2 μg/mL heparin, 150 μg/mL BSA, GlutaMAX (Life Technologies), N2 supplement (Life Technologies), B27 supplement (Life Technologies), 10 ng/mL EGF (Life Technologies), 10 ng/mL FGF (Life Technologies), as previously described(Pollard et al. 2009; MacLeod et al. 2019). Cell dissociation was performed using Accutase (Sigma Aldrich). All cell lines were maintained at 37 °C with 5% CO2. HEK293T cells were cultured in DMEM media supplemented with 10% FBS and 1% pen strep(Life Technologies). Where needed, puromycin was added to the media at a concentration of 2 μg/mL. The following drugs, when used, were dissolved in DMSO: A-485 (Tocris, Cat. No. 6387), and Temozolomide (MilliporeSigma).

All GSCs were confirmed to match their parental primary GBM tumors through STR profiling at The Centre for Applied Genomics (Toronto, Ontario, Canada). Cells were periodically subjected to testing with the MycoAltert Plus mycoplasma detection kit (Lonza Biosciences) to confirm the absence of contamination.

### Genome-wide CRISPR-Cas9 Knock out screen

CRISPR-Cas9 KO screen was performed as previously described using the 70K TKOv3 library (Addgene Pooled Library #90294)(MacLeod et al. 2019; Hart et al. 2017). 4×10^8^ cells were transduced with M.O.I (Multiplicity of infection) 0.3 in presence of 0.8 ug/mL of polybrene and plated into 10cm plates. Twenty-four hours post virus transduction, media was removed and fresh GSC media containing 2 μg/mL puromycin was added to cells. After three days of puromycin selection, cells were harvested, counted and 2×10^7^ cells were collected as T0 samples and stored at −80°C for later processing. The remaining pool of cells were divided equally into three biological replicates, plus one “unsorted” replicate. In the entire process, a minimum 200-fold library coverage was maintained for all replicates. After 4 additional days of culture, cells were harvested using Accutase. The three sorted replicate were washed once with PBS, blocked on ice for 30 with BSA 3%, and stained with CD44 antibody (1:200, Invitrogen Cat No: 56044180) and eBioscience Fixable Viability Dye eFluor 450 (1:500, Invitrogen Cat No:65-0863-14) for 1 hr on ice in the dark. After staining, cells were washed 2X with PBS and fixed 1% Paraformaldehyde (PFA) for 10 minutes at room temperature in the dark. Cells were washed with PBS and resuspended in FACS buffer (PBS, 0.5% BSA, 1mM EDTA) and stored at 4°C. FACS was performed on BD Influx and BD FACS Aria III instruments (BD Biosciences). The top 15% and bottom 15% of CD44-expressing cells per replicate were collected and the cell pellets were frozen at -80°C for further processing.

### Genomic DNA Library Preparation and Sequencing

Genomic DNA extraction from frozen cell pellets was performed as previously described(F. Chen et al. 2011). For each sample, 50 μg DNA was processed for illumina sequencing using a unique combination of i5 and i7 barcodes as previously described (MacLeod et al. 2019; MacLeod, Rajakulendran, and Angers 2022). The barcoded PCR products were then gel purified and sequenced with NextSeq500 instrument (Illumina) at a read depth of 200-fold for all samples. Next generation sequencing was performed at the Network Biology Collaborative Centre, Lunenfeld-Tanenbaum Research Institute (nbcc.lunenfeld.ca, Toronto).

### CRISPR-Cas9 gene editing

To generate single gene knockout, individual gRNAs were ligated into Esp3I digested LentiCRISPRV2 (LCV2) vector (Addgene catalog #52961). GSCs were infected with LCV2 for gene knockout by transducing 3×10^5^ cells in the presence of 0.8 μg/mL polybrene. After 24hr, cells were selected with 2 μg/mL puromycin for 3 days prior to seeding for performing various experiments. To quantify gene editing efficiency, TIDE (Brinkman et al. 2014) was used from PCR amplicons flanking gRNA target sites. For a complete list of gRNA sequences, see Table S2.

### Lentivirus production

The lentivirus containing the TKOv3 gRNA library was prepared as described previously (Hart et al. 2017). For small scale lentiviral production for single gRNA, 3.5×10^6^ HEK293T cells were seeded on 10 cm plates and the next day transfected with 6 μg lentiviral plasmid, 6 μg of psPAX2 and 2 μg of pMD2.G (psPAX2: Addgene #12260 and pMD2.G: Addgene #12259). Media was replaced 24 hours post-transfection and viral media containing the virus was harvested 48 hours post-transfection. Viral supernatant was then centrifuged at 500xg for 3 minutes, filtered through a Stericup-HV PVDF 0.45-μm filter, and concentrated using the Lenti-X Concentrator solution (Clontech) according to manufacturer’s protocol.

### RNA sequencing

Cells were transduced with LCV2 carrying gRNAs targeting *EP300* or *AAVS1* and seeded on coated 6-well plates. After 24hr, media was replaced with fresh media containing 2 μg/mL puromycin. After 1 week, total RNA was isolated from cells using the TRIzol reagent (Thermofisher). The samples were submitted to the Donnelly Sequencing Center at the University of Toronto (https://thedonnellycentre.utoronto.ca/donnelly-sequencing-centre) for sequencing. The quality of RNA was measured using BioAnalyzer (Agilent) and RNA-seq libraries (4 biological replicates each sample) were derived using NEBNext Poly(A) mRNA Magnetic isolation module followed by NEBNext Ultra II Directional RNA library prep kit. Libraries were loaded on an NovaSeq6000 (Illumina) at 320 pMol with 150 paired-end reads. The trimmed reads were aligned to the reference genome (UCSC hg19) using Kallisto(Bray et al. 2016) and differential expression analysis was performed using the DESeq2 package(Love, Huber, and Anders 2014). Pathway, gene ontology and GSEA analyses were performed using gProfiler and GSEA software (Subramanian et al. 2005; Reimand et al. 2007).

### Quantitative Polymerase Chain Reaction (qPCR)

RNA was extracted using the TRIzol reagent (Thermofisher) and 2 ug of total RNA was used for cDNA synthesis using Super-Script II reverse transcription kit (Thermo Fisher). qPCR was performed using SyBrGreen (ThermoFisher) on a BioRAD CFX instrument. The primer sequences can be found in Table S3.

### Cell proliferation assay

Cells with *AAVS1* gRNA and *EP300* KO were plated on 6 well plates (4 days after transduction and puro selection) and were imaged using the Incucyte live cell imaging platform (Sartorius). Quantification of cell confluency from images was performed using Incucyte instrument software (version 2023A Rev2).

### In vitro Limiting Dilution Assay

*In vitro* sphere-forming ability was measured by culturing GSCs in serial dilutions (range of 2–1000 cells per well) on non-adherent 96-well plates. After 2-3 weeks of culture, the frequency of sphere forming cells was quantified by inequality in frequency between multiple groups and tested for adequacy of the single-hit model using Extreme limiting dilution analysis (ELDA) software (http://bioinf.wehi.edu.au/software/elda) (Hu and Smyth 2009).

### TMZ cell viability assay

GSCs were transduced with LCV2 *AAVS1* and *EP300*, and after 24hr, media was replaced with fresh media containing puromycin. After selection, 2000 cells/well were seeded in 96 well plates. Twenty-four hours after seeding cells were treated with TMZ. The dose-response experiments in these cells were conducted in the range of 0-100 µM of TMZ for 7 days with one media change with new drug added at mid-point. Relative cell viability was assessed using the *CellTiter*-*Glo* Luminescent Cell Viability Assay (Promega).

### Flow cytometry

GSCs were harvested from plates using accutase and then washed with cold PBS. For assessment of CD44 surface expression, blocking of cells in 3% BSA/PBS (blocking buffer) was performed for 30 minutes on ice, followed by primary antibody (anti-CD44 Alexa Fluor 700, Invitrogen Cat no: 56044180) at 1:200, and viability dye (Fixable Viability Dye eFluor™ 450, Invitrogen Cat No:65-0863-14) at 1:500, in blocking buffer for 1 hour on ice in darkness. Subsequently, cells were washed 2x with PBS and fixed in 1% PFA for 10 minutes followed by resuspension in PBS for flow cytometry. CytoFlex S (Beckman Coulter) instrument and the CytExpert software was used for flow cytometry and data was analyzed and visualized using FlowJo™ v10.8 Software (BD Life Sciences).

### Invasion assay

Hylarounic acid based Diels–Alder click-cross-linked hydrogel was prepared as described previously(Smith et al. 2023). For the invasion assay, 15 µL of hydrogel solution was plated into each well of a 384-well plate. Following gelation, hydrogels were washed extensively: with PBS, NS-A basal media, and with GSC growth medium with at least 45 min incubation between wash steps. Prior to seeding cells on hydrogels, GSCs were either transduced with LCV2 viruses (*AAVS1* and *EP300*) or treated with DMSO and A-485. After one week of KO/pre-treatment, GSC lines were plated on top hydrogels at a density of 3500 cells in 60 μL of media per hydrogel. 24 hours after seeding, another 15 μL of fresh media was added to the cells. Four days after plating, cells were fixed with 4% PFA for 45 min at room temperature and counterstained with Hoechst (1:500 Invitrogen H1399) to visualize nuclei. Approximately 10 000 red fluorescent beads (1:15 Fluospheres 15 μm polystyrene beads, Thermo Fisher Scientific F8842) were then added to mark the surface of the hydrogel and allowed to settle for 10 min prior to imaging. Hydrogels were imaged using an Olympus confocal microscope (FV1000), taking images every 20 μm on the *z*-axis. Images were analyzed using Imaris 8.3.1 (Bitplane) software and processed using MATLAB(Smith et al. 2023).

### Nuclei isolation and cleavage under targets and release using nuclease (CUT&RUN)

Cells were transduced with LCV2 *EP300* or *AAVS1* and seeded on coated 6-well plates. After 24hr, media was replaced with fresh media containing 2 μg/mL puromycin. After 1 week, cells were harvested for CUT&RUN assay. To crosslink proteins to the genomic DNA, the final concentration of 0.1% formaldehyde was added to 500,000 cells in GBM media and incubated for 1 minute. Then, Glycine with a final concentration of 125mM was added to cells to quench the crosslink. CUT&RUN was performed using the CUTANA ChIC/CUT&RUN kit (14-1048, EpiCypher). Briefly, after crosslinking, cells were spin down at 600xg for 3 minutes and washed 2x with the kit wash buffer. Meanwhile, concanavalin A beads (ConA, 14–1048, EpiCypher) were activated using beads activation buffer (14–1048, EpiCypher). Then crosslinked cells were added to the antibody buffer. Antibodies were added and incubated overnight at 4 degrees on the nutator. Next day, cells were washed with the permeabilization buffer and CUTANA pAG-MNase was added to the cells. Also, 100mM CaCl_2_ was added to activate MNase to cleave target chromatin and cells were incubated at 4 degree. After 2 hours, CUTANA stop buffer was added to the extracted chromatin. The crosslink was reversed by adding 10% PFA and 20mg/mL protease to the extracted chromatin and incubation at 55℃ overnight. The next day, DNA was purified using the CUTANA ChIC/CUT&RUN kit. The samples were submitted to the Donnelly Sequencing Center, at the University of Toronto for sequencing. Libraries were prepared using NEBNext Ultra II FS DNA library prep kit for illumina and quantified using the Qubit 1X dsDNA high sensitivity assay. Sizing was done using Agilent TapeStation D1000 ScreenTape and sequencing was loaded on the NovaSeq6000 (Illumina) at 300 pMol concentration with 150 paired-end reads. H3K27Ac and p300 antibodies were as followed respectively: H3K27Ac Monoclonal Antibody, Invitrogen (MA5-23516), p300 (D8Z4E) Rabbit mAb, cell signaling (product #86377).

### In vivo study

Animal husbandry, ethical handling of mice and all animal work were carried out according to guidelines approved by Canadian Council on Animal Care and under protocols approved by the Toronto Centre for Phenogenomics Animal Care Committee (18-0272H). R26-LSL-Cas9-GFP (JAX #026175), Trp53^fl/fl^ (Trp53tm1Brn, JAX #008462), Pten^fl/fl^ (Ptentm1Hwu/J, JAX #006440), Rb1^fl/f l^ (Rb1tm2Brn/J, JAX #026563) were animals used in this study and they were all obtained from the Jackson laboratories. An equal number of male and female animals without any bias.

### Lentivirus production for *in vivo* study

gRNAs targeting mouse *Ep300* (5’-AGCAAGCTAATGGGGAAGTGAGG-3’, 5’-AGGAACTAGAAGAGAAACGAAGG-3’ ) were cloned into pLKO-Cre stuffer v4 plasmid (Addgene #158032) by using BsmBI restriction sites. 293T cells (Invitrogen R700-07) were seeded 10 cm plates and transfected for 8 hours using PEI (polyethyleneimine) in non-serum media conditions with gRNA containing lentiviral construct, packaging plasmids psPAX2 and pPMD2.G. Post transfection, media was added to the plates supplemented with 10% Fetal bovine serum and 1% Pencillin-Streptomycin antibiotic solution (w/v). 48 hours after transfection, viral supernatant was filtered through a Stericup-HV PVDF 0.45-μm filter, and then concentrated ∼2,000-fold by ultracentrifugation in an MLS-50 rotor (Beckman Coulter). Viral titers were calculated by flow cytometry-based quantification of infected R26-LSL-tdTomato MEFs.

### Intracranial injection and *in vivo* lentiviral transduction

Concentrated virus was mixed with 0.05% Fast Green (F7252-5G) and loaded into a syringe (Hamilton 7659-01) with 33-gauge needle (Hamilton 7803-05). P0 pups were anesthetized on parafilm covered ice and their head secured with a custom 3D-printed mold. A stereotactic manipulator was used to position the needle to 0.3 mm above the Bregma towards the Lambda Suture and 0.1 mm lateral of Sagittal Suture into the right ventricle. The needle punctured 3 mm into the skull, and retracted 1mm for a final depth of 2 mm. 1 µL of virus was administered and allowed 1 minute to diffuse before retraction of the needle. Post-injection, the neonates were warmed on a heating pad.

### Quantification and statistical analysis

#### Analysis of Genome-wide CRISPR-Cas9 FACS-based screen

Raw sequencing data was demultiplexed and trimmed the adaptor sequence from FASTQ files prior to mapping of sequencing reads to the TKOv3 library using the MAGeCK algorithm (W. Li et al. 2014) *count* function. Gene knockouts positively selected in the CD44 low population bin over the CD44 high population bin were identified using the *test* function in MaGeCK (FDR < 0.2).

### Analysis of CUT&RUN

We used the modified pipeline developed by Henikoff lab (Zheng, Ahmad, and Henikoff 2020) to align the fastq files into hg38 genome and process them into Sam, Bam, bigwig and bedgraph files. For peakCalling, we used SEACR (Sparse Enrichment Analysis for CUT&RUN) with cut off 0.01 for H3K27Ac peak calls and 0.001 for P300 binding sites call. Homer software was used to annotate the peaks (annotatePeaks.pl) for H3K27Ac peaks and p300 binding sites (Heinz et al. 2010). Enhancers were identified using the ROSE algorithm on H3K27Ac samples (Whyte et al. 2013). Enhancer regions in each sample were tested against all samples to identify the significantly enriched motifs (Heinz et al. 2010). P300 bound regions were also tested with Homer software (findMotifsGenome.pl) to find the enrichment of TF binding motifs(Heinz et al. 2010). To identify IR motifs in ATAC-seq, we generated a consensus file for IR GSCs peaks, and also one master consensus peak file containing both IR and Dev GSCs open regions, then we tested the IR consensus peak regions against the master peak file(Guilhamon et al. 2021).

### Statistical analysis

The details related to the statistical analysis for each experiment can be found in figure legends. The Graphpad Prism 5 (GraphPad Software, La Jolla, California, USA) was used for statistical analysis, unless specified otherwise. All statistical tests were two-sided unless specified otherwise. Statistical analysis for flow cytometry, and invasion assay was performed using unpaired, two-sample t tests from n = 3 replicates. Statistical analysis of qPCR data was performed using GraphPad Prism and two-way ANOVA test was used to calculate significance. Values were obtained from at least 2 biological replicates for each sample and hcyclo was used for normalization. In vitro LDA was performed in ELDA software (see above) and SFC is plotted as 95% confidence intervals for 1/stem cell frequency. To assess the differences in stem cell frequencies between various groups Pairwise chi-square tests were used.

## Data availability

RNA-Seq data generated from this study is deposited in GEO (GSE290575). CUT&RUN sequencing data and associated peak files are also available in GEO (GSE290646). All other raw data generated in this study are available upon request from the corresponding author.

## Acknowledgements

The authors of this study wish to thank Sherin Mohammed Shibin at the Donnelly Sequencing Centre, (http://ccbr.utoronto.ca/donnelly-sequencing-centre, Toronto) and Kin Chan at the Network Biology Collaborative Centre (nbcc.lunenfeld.ca, Toronto) for next-generation sequencing services. We also want to thank all members of Angers lab for their valuable discussion throughout the course of this study and members of the Dirks lab for their assistance in providing patient-derived GSCs using in this study. Also we would like to thank Dr. Lupien group specially Ankita Nand for their valuable advice and guidance for CUT&RUN analysis. This research was supported by the Canadian Institutes of Health Research (PJT169054 to S. Angers).

## References

Bhat, Krishna P. L., Veerakumar Balasubramaniyan, Brian Vaillant, Ravesanker Ezhilarasan, Karlijn Hummelink, Faith Hollingsworth, Khalida Wani, et al. 2013. “Mesenchymal Differentiation Mediated by NF-κB Promotes Radiation Resistance in Glioblastoma.” Cancer Cell 24 (3): 331–46.

Bhat, Krishna P. L., Katrina L. Salazar, Veerakumar Balasubramaniyan, Khalida Wani, Lindsey Heathcock, Faith Hollingsworth, Johanna D. James, et al. 2011. “The Transcriptional Coactivator TAZ Regulates Mesenchymal Differentiation in Malignant Glioma.” Genes & Development 25 (24): 2594–2609.

Bray, Nicolas L., Harold Pimentel, Páll Melsted, and Lior Pachter. 2016. “Near-Optimal Probabilistic RNA-Seq Quantification.” Nature Biotechnology 34 (5): 525–27.

Brinkman, Eva K., Tao Chen, Mario Amendola, and Bas van Steensel. 2014. “Easy Quantitative Assessment of Genome Editing by Sequence Trace Decomposition.” Nucleic Acids Research 42 (22): e168.

Chanoch-Myers, Rony, Adi Wider, Mario L. Suva, and Itay Tirosh. 2022. “Elucidating the Diversity of Malignant Mesenchymal States in Glioblastoma by Integrative Analysis.” Genome Medicine 14 (1): 106.

Chen, Fuqiang, Shondra M. Pruett-Miller, Yuping Huang, Monika Gjoka, Katarzyna Duda, Jack Taunton, Trevor N. Collingwood, Morten Frodin, and Gregory D. Davis. 2011. “High-Frequency Genome Editing Using ssDNA Oligonucleotides with Zinc-Finger Nucleases.” Nature Methods 8 (9): 753–55.

Chen, Qingjuan, Binhui Yang, Xiaochen Liu, Xu D. Zhang, Lirong Zhang, and Tao Liu. 2022. “Histone Acetyltransferases CBP/p300 in Tumorigenesis and CBP/p300 Inhibitors as Promising Novel Anticancer Agents.” Theranostics 12 (11): 4935–48.

Chow, Lionel M. L., Raelene Endersby, Xiaoyan Zhu, Sherri Rankin, Chunxu Qu, Junyuan Zhang, Alberto Broniscer, David W. Ellison, and Suzanne J. Baker. 2011. “Cooperativity within and among Pten, p53, and Rb Pathways Induces High-Grade Astrocytoma in Adult Brain.” Cancer Cell 19 (3): 305–16.

Dancy, Beverley M., and Philip A. Cole. 2015. “Protein Lysine Acetylation by p300/CBP.” Chemical Reviews 115 (6): 2419–52.

De Bono, Johann S., Elena Cojocaru, Elizabeth Ruth Plummer, Tomasz Knurowski, Karen Clegg, Fay Ashby, Neil Pegg, William West, and Anthony Nigel Brooks. 2019. “An Open Label Phase I/IIa Study to Evaluate the Safety and Efficacy of CCS1477 as Monotherapy and in Combination in Patients with Advanced Solid/metastatic Tumors.” Journal of Clinical Oncology: Official Journal of the American Society of Clinical Oncology 37 (15_suppl): TPS5089–TPS5089.

Dirkse, Anne, Anna Golebiewska, Thomas Buder, Petr V. Nazarov, Arnaud Muller, Suresh Poovathingal, Nicolaas H. C. Brons, et al. 2019. “Stem Cell-Associated Heterogeneity in Glioblastoma Results from Intrinsic Tumor Plasticity Shaped by the Microenvironment.” Nature Communications 10 (1): 1–16.

Fan, Xiaoqing, Junqi Fan, Haoran Yang, Chenggang Zhao, Wanxiang Niu, Zhiyou Fang, and Xueran Chen. 2021. “Heterogeneity of Subsets in Glioblastoma Mediated by Smad3 Palmitoylation.” Oncogenesis 10 (10): 72.

Guilhamon, Paul, Charles Chesnelong, Michelle M. Kushida, Ana Nikolic, Divya Singhal, Graham MacLeod, Seyed Ali Madani Tonekaboni, et al. 2021. “Single-Cell Chromatin Accessibility Profiling of Glioblastoma Identifies an Invasive Cancer Stem Cell Population Associated with Lower Survival.” eLife 10 (January). 10.7554/eLife.64090.

Hara, Toshiro, Rony Chanoch-Myers, Nathan D. Mathewson, Chad Myskiw, Lyla Atta, Lillian Bussema, Stephen W. Eichhorn, et al. 2021. “Interactions between Cancer Cells and Immune Cells Drive Transitions to Mesenchymal-like States in Glioblastoma.” Cancer Cell 39 (6): 779–92.e11.

Hart, Traver, Amy Hin Yan Tong, Katie Chan, Jolanda Van Leeuwen, Ashwin Seetharaman, Michael Aregger, Megha Chandrashekhar, et al. 2017. “Evaluation and Design of Genome-Wide CRISPR/SpCas9 Knockout Screens.” G3 7 (8): 2719–27.

Heinz, Sven, Christopher Benner, Nathanael Spann, Eric Bertolino, Yin C. Lin, Peter Laslo, Jason X. Cheng, Cornelis Murre, Harinder Singh, and Christopher K. Glass. 2010. “Simple Combinations of Lineage-Determining Transcription Factors Prime Cis-Regulatory Elements Required for Macrophage and B Cell Identities.” Molecular Cell 38 (4): 576–89.

Hu, Yifang, and Gordon K. Smyth. 2009. “ELDA: Extreme Limiting Dilution Analysis for Comparing Depleted and Enriched Populations in Stem Cell and Other Assays.” Journal of Immunological Methods 347 (1-2): 70–78.

Khan, A. Basit, Sungho Lee, Akdes Serin Harmanci, Rajan Patel, Khatri Latha, Yuhui Yang, Anantha Marisetty, et al. 2023. “CXCR4 Expression Is Associated with Proneural-to-Mesenchymal Transition in Glioblastoma.” International Journal of Cancer. Journal International Du Cancer 152 (4): 713–24.

Lan, Xiaoyang, David J. Jörg, Florence M. G. Cavalli, Laura M. Richards, Long V. Nguyen, Robert J. Vanner, Paul Guilhamon, et al. 2017. “Fate Mapping of Human Glioblastoma Reveals an Invariant Stem Cell Hierarchy.” Nature 549 (7671): 227–32.

Lasko, Loren M., Clarissa G. Jakob, Rohinton P. Edalji, Wei Qiu, Debra Montgomery, Enrico L. Digiammarino, T. Matt Hansen, et al. 2017. “Discovery of a Selective Catalytic p300/CBP Inhibitor That Targets Lineage-Specific Tumours.” Nature 550 (7674): 128–32.

Lau, Jasmine, Shirin Ilkhanizadeh, Susan Wang, Yekaterina A. Miroshnikova, Nicolas A. Salvatierra, Robyn A. Wong, Christin Schmidt, Valerie M. Weaver, William A. Weiss, and Anders I. Persson. 2015. “STAT3 Blockade Inhibits Radiation-Induced Malignant Progression in Glioma.” Cancer Research 75 (20): 4302–11.

Li, Chien-Hsiu, Chih-Yeu Fang, Ming-Hsien Chan, Pei-Jung Lu, Luo-Ping Ger, Jan-Show Chu, Yu-Chan Chang, Chi-Long Chen, and Michael Hsiao. 2023. “The Activation of EP300 by F11R Leads to EMT and Acts as a Prognostic Factor in Triple-Negative Breast Cancers.” Hip International: The Journal of Clinical and Experimental Research on Hip Pathology and Therapy 9 (3): 165–81.

Li, Wei, Han Xu, Tengfei Xiao, Le Cong, Michael I. Love, Feng Zhang, Rafael A. Irizarry, Jun S. Liu, Myles Brown, and X. Shirley Liu. 2014. “MAGeCK Enables Robust Identification of Essential Genes from Genome-Scale CRISPR/Cas9 Knockout Screens.” Genome Biology 15 (12): 554.

Louis, David N., Arie Perry, Pieter Wesseling, Daniel J. Brat, Ian A. Cree, Dominique Figarella-Branger, Cynthia Hawkins, et al. 2021. “The 2021 WHO Classification of Tumors of the Central Nervous System: A Summary.” Neuro-Oncology 23 (8): 1231–51.

Love, Michael I., Wolfgang Huber, and Simon Anders. 2014. “Moderated Estimation of Fold Change and Dispersion for RNA-Seq Data with DESeq2.” Genome Biology 15 (12): 550.

Lv, Xuejiao, Qian Li, Hang Liu, Meihan Gong, Yingying Zhao, Jinyang Hu, Fan Wu, and Xudong Wu. 2022. “JUN Activation Modulates Chromatin Accessibility to Drive TNFα-Induced Mesenchymal Transition in Glioblastoma.” Journal of Cellular and Molecular Medicine 26 (16): 4602–12.

Ma, Chaoqun, Shuhong Huang, Lei Xu, Li Tian, Yan Yang, and Jianming Wang. 2020. “Transcription Co-Activator P300 Activates Elk1-aPKC-ι Signaling Mediated Epithelial-to-Mesenchymal Transition and Malignancy in Hepatocellular Carcinoma.” Oncogenesis 9 (3): 32.

MacLeod, Graham, Danielle A. Bozek, Nishani Rajakulendran, Vernon Monteiro, Moloud Ahmadi, Zachary Steinhart, Michelle M. Kushida, et al. 2019. “Genome-Wide CRISPR-Cas9 Screens Expose Genetic Vulnerabilities and Mechanisms of Temozolomide Sensitivity in Glioblastoma Stem Cells.” Cell Reports 27 (3): 971–86.e9.

MacLeod, Graham, Fatemeh Molaei, Shahan Haider, Maira P. Almeida, Sichun Lin, Michelle Kushida, Haresh Sureshkumar, et al. 2024. “Fitness Screens Map State-Specific Glioblastoma Stem Cell Vulnerabilities.” Cancer Research, August. 10.1158/0008-5472.CAN-23-4024.

MacLeod, Graham, Nishani Rajakulendran, and Stephane Angers. 2022. “Identification of Drug Resistance Mechanisms Using Genome-Wide CRISPR-Cas9 Screens.” Methods in Molecular Biology (Clifton, N.J.) 2535:141–56.

Marques, Carolina, Thomas Unterkircher, Paula Kroon, Barbara Oldrini, Annalisa Izzo, Yuliia Dramaretska, Roberto Ferrarese, et al. 2021. “NF1 Regulates Mesenchymal Glioblastoma Plasticity and Aggressiveness through the AP-1 Transcription Factor FOSL1.” eLife 10 (August). 10.7554/eLife.64846.

Matusow, Bernice, W. Spevak, Chao Zhang, Yan Ma, Rafe Shellooe, J. Tsai, Peipei Li, et al. 2024. “Abstract 660: OPN-6602, a Potent Dual EP300/CBP Bromodomain Inhibitor, Targets Multiple Myeloma through Concomitant Suppression of IRF4 and c-MYC.” *Cancer Research*, March. 10.1158/1538-7445.am2024-660.

Muthukrishnan, Sree Deepthi, Riki Kawaguchi, Pooja Nair, Rachna Prasad, Yue Qin, Maverick Johnson, Qing Wang, et al. 2022. “P300 Promotes Tumor Recurrence by Regulating Radiation-Induced Conversion of Glioma Stem Cells to Vascular-like Cells.” Nature Communications 13 (1): 6202.

Neftel, Cyril, Julie Laffy, Mariella G. Filbin, Toshiro Hara, Marni E. Shore, Gilbert J. Rahme, Alyssa R. Richman, et al. 2019. “An Integrative Model of Cellular States, Plasticity, and Genetics for Glioblastoma.” Cell 178 (4): 835–49.e21.

Nicosia, Luciano, Gary J. Spencer, Nigel Brooks, Fabio M. R. Amaral, Naseer J. Basma, John A. Chadwick, Bradley Revell, et al. 2023. “Therapeutic Targeting of EP300/CBP by Bromodomain Inhibition in Hematologic Malignancies.” Cancer Cell 41 (12): 2136–53.e13.

Ogryzko, Vasily V., R. Louis Schiltz, Valya Russanova, Bruce H. Howard, and Yoshihiro Nakatani. 1996. “The Transcriptional Coactivators p300 and CBP Are Histone Acetyltransferases.” Cell 87 (5): 953–59.

Phillips, Heidi S., Samir Kharbanda, Ruihuan Chen, William F. Forrest, Robert H. Soriano, Thomas D. Wu, Anjan Misra, et al. 2006. “Molecular Subclasses of High-Grade Glioma Predict Prognosis, Delineate a Pattern of Disease Progression, and Resemble Stages in Neurogenesis.” Cancer Cell 9 (3): 157–73.

Pollard, Steven M., Koichi Yoshikawa, Ian D. Clarke, Davide Danovi, Stefan Stricker, Roslin Russell, Jane Bayani, et al. 2009. “Glioma Stem Cell Lines Expanded in Adherent Culture Have Tumor-Specific Phenotypes and Are Suitable for Chemical and Genetic Screens.” Cell Stem Cell 4 (6): 568–80.

Rajakulendran, Nishani, Katherine J. Rowland, Hayden J. Selvadurai, Moloud Ahmadi, Nicole I. Park, Sergey Naumenko, Sonam Dolma, et al. 2019. “Wnt and Notch Signaling Govern Self-Renewal and Differentiation in a Subset of Human Glioblastoma Stem Cells.” Genes & Development 33 (9-10): 498–510.

Reik, W., W. Dean, and J. Walter. 2001. “Epigenetic Reprogramming in Mammalian Development.” Science 293 (5532): 1089–93.

Reimand, Jüri, Meelis Kull, Hedi Peterson, Jaanus Hansen, and Jaak Vilo. 2007. “g:Profiler--a Web-Based Toolset for Functional Profiling of Gene Lists from Large-Scale Experiments.” Nucleic Acids Research 35 (Web Server issue): W193–200.

Richards, Laura M., Owen K. N. Whitley, Graham MacLeod, Florence M. G. Cavalli, Fiona J. Coutinho, Julia E. Jaramillo, Nataliia Svergun, et al. 2021. “Gradient of Developmental and Injury Response Transcriptional States Defines Functional Vulnerabilities Underpinning Glioblastoma Heterogeneity.” Nature Cancer 2 (2): 157–73.

Schmitt, Matthias Jürgen, Company, Carlos, Yuliia Dramaretska, Iros Barozzi, Andreas Göhrig, Sonia Kertalli, Melanie Großmann, et al. 2021. “Phenotypic Mapping of Pathologic Cross-Talk between Glioblastoma and Innate Immune Cells by Synthetic Genetic Tracing.” Cancer Discovery 11 (3): 754–77.

Segerman, Anna, Mia Niklasson, Caroline Haglund, Tobias Bergström, Malin Jarvius, Yuan Xie, Ann Westermark, et al. 2016. “Clonal Variation in Drug and Radiation Response among Glioma-Initiating Cells Is Linked to Proneural-Mesenchymal Transition.” Cell Reports 17 (11): 2994–3009.

Singh, Sheila K., Cynthia Hawkins, Ian D. Clarke, Jeremy A. Squire, Jane Bayani, Takuichiro Hide, R. Mark Henkelman, Michael D. Cusimano, and Peter B. Dirks. 2004. “Identification of Human Brain Tumour Initiating Cells.” Nature 432 (7015): 396–401.

Smith, Laura J., Arianna Skirzynska, Allysia A. Chin, Amy E. Arnold, Michelle Kushida, Peter B. Dirks, and Molly S. Shoichet. 2023. “Engineered In Vitro Tumor Model Recapitulates Molecular Signatures of Invasion in Glioblastoma.” ACS Materials Au 3 (5): 514–27.

Subramanian, Aravind, Pablo Tamayo, Vamsi K. Mootha, Sayan Mukherjee, Benjamin L. Ebert, Michael A. Gillette, Amanda Paulovich, et al. 2005. “Gene Set Enrichment Analysis: A Knowledge-Based Approach for Interpreting Genome-Wide Expression Profiles.” Proceedings of the National Academy of Sciences of the United States of America 102 (43): 15545–50.

Suvà, Mario L., Nicolo Riggi, and Bradley E. Bernstein. 2013. “Epigenetic Reprogramming in Cancer.” Science 339 (6127): 1567–70.

Thornton, Nicole, Vanja Karamatic Crew, Louise Tilley, Carole A. Green, Chwen Ling Tay, Rebecca E. Griffiths, Belinda K. Singleton, et al. 2020. “Disruption of the Tumour-Associated EMP3 Enhances Erythroid Proliferation and Causes the MAM-Negative Phenotype.” Nature Communications 11 (1): 3569.

Wang, Qianghu, Baoli Hu, Xin Hu, Hoon Kim, Massimo Squatrito, Lisa Scarpace, Ana C. deCarvalho, et al. 2017. “Tumor Evolution of Glioma-Intrinsic Gene Expression Subtypes Associates with Immunological Changes in the Microenvironment.” Cancer Cell 32 (1): 42–56.e6.

Whyte, Warren A., David A. Orlando, Denes Hnisz, Brian J. Abraham, Charles Y. Lin, Michael H. Kagey, Peter B. Rahl, Tong Ihn Lee, and Richard A. Young. 2013. “Master Transcription Factors and Mediator Establish Super-Enhancers at Key Cell Identity Genes.” Cell 153 (2): 307–19.

Zhang, Dandan, Yanshuang Peng, Gaoren Lin, Qian Xiao, Han Liu, Xiaoxu Nan, Jinpeng Han, et al. 2023. “TGF-βRII/EP300/SMAD4 Cascade Signaling Pathway Promotes Invasion and Glycolysis in Oral Squamous Cell Carcinoma.” Journal of Oral Pathology & Medicine: Official Publication of the International Association of Oral Pathologists and the American Academy of Oral Pathology 52 (6): 483–92.

Zhang, Junchang, Han Wang, Jing Wu, Cheng Yuan, Songyao Chen, Shuhao Liu, Mingyu Huo, Changhua Zhang, and Yulong He. 2022. “GALNT1 Enhances Malignant Phenotype of Gastric Cancer via Modulating CD44 Glycosylation to Activate the Wnt/β-Catenin Signaling Pathway.” International Journal of Biological Sciences 18 (16): 6068–83.

Zheng, Ye, Kami Ahmad, and Steven Henikoff. 2020. “CUT&Tag Data Processing and Analysis Tutorial.” https://www.protocols.io/view/cut-amp-tag-data-processing-and-analysis-tutorial-bjk2kkye.html.

